# Simulation of visual perception and learning with a retinal prosthesis

**DOI:** 10.1101/206409

**Authors:** James R. Golden, Cordelia Erickson-Davis, Nicolas P. Cottaris, Nikhil Parthasarathy, Fred Rieke, David H. Brainard, Brian A. Wandell, E.J. Chichilnisky

## Abstract

The nature of artificial vision with a retinal prosthesis, and the degree to which the brain can adapt to the unnatural input from such a device, are poorly understood. Therefore, the development of current and future devices may be aided by theory and simulations that help to infer and understand what prosthesis patients see. A biologically-informed, extensible computational framework is presented here to predict visual perception and the potential effect of learning with a subretinal prosthesis. The framework relies on optimal linear reconstruction of the stimulus from retinal responses to infer the visual information available to the patient. A simulation of the physiological optics of the eye and light responses of the major retinal neurons was used to calculate the optimal linear transformation for reconstructing natural images from retinal activity. The result was then used to reconstruct the visual stimulus during the artificial activation expected from a subretinal prosthesis in a degenerated retina, as a proxy for inferred visual perception. Several simple observations reveal the potential utility of such a simulation framework. The inferred perception obtained with prosthesis activation was substantially degraded compared to the inferred perception obtained with normal retinal responses, as expected given the limited resolution and lack of cell type specificity of the prosthesis. Consistent with clinical findings and the importance of cell type specificity, reconstruction using only ON cells, and not OFF cells, was substantially more accurate. Finally, when reconstruction was re-optimized for prosthesis stimulation, simulating the greatest potential for learning by the patient, the accuracy of inferred perception was much closer to that of healthy vision. The reconstruction approach thus provides a more complete method for exploring the potential for treating blindness with retinal prostheses than has been available previously. It may also be useful for interpreting patient data in clinical trials, and for improving prosthesis design.

## Introduction

Retinal prostheses can restore visual perception to blind patients through electrical stimulation of surviving neurons in the retina, based on images captured by a camera (Humayun et al. 2012; Zrenner et al. 2011). The perceptual experience of patients with existing devices, however, is far from that of a person with normal vision. Several efforts are underway to build next generation devices (Mathieson et al. 2012; Lorach et al. 2015a). Furthermore, clinical experience with other types of neural implants suggests that learning by the patient may help overcome some device limitations (Hallberg and Ringdahl 2004; Shannon 2012; Eisen 2003; Waltzman et al. 1993). Nonetheless, the degree to which technical improvements and learning can in principle improve outcomes remains poorly understood, and current understanding of the experience of artificial vision itself is limited. Thus, it would be valuable to have a computational framework to predict the visual experience that retinal prostheses provide to patients, as well as how that visual experience could potentially improve with learning (Beyeler et al. 2017). Such a framework could help researchers and clinicians interpret what patients see and how they learn, and could guide future technical development by identifying factors that limit performance.

The goal of this work is to provide a starting point and proof of concept for such a framework. To do this, we use optimal reconstruction of the incident stimulus from the simulated retinal output to infer the information about the stimulus available to the central visual system. Although reconstruction of a sensory stimulus from neural signals has been used in previous work (Rieke 1997, Warland 1997), it has not been applied to inferring visual perception and learning with a prosthesis (but see Nirenberg and Pandarinath 2012). The reconstruction framework approach has the advantage of providing a direct assessment of the visual sensations potentially experienced by the patient, rather than an indirect representation in terms of patterns of electrical activity in the optic nerve. Unlike existing approaches (Beyeler et al. 2017), it has a theoretical foundation, and in principle it can be extended from the starting point presented here to include all known aspects of retinal signals and retinal prostheses. Finally, the approach provides a straightforward way to reason about how a patient with a prosthesis could potentially learn to use it effectively, a fundamental aspect of neural interfaces for which few quantitative reasoning tools are available.

Here we develop the reconstruction approach, and provide an extensible software framework implementing it. The framework is based on simple simulations of healthy retinal function (<u>ISETBio</u>; Cottaris et al. 2018; Jiang et al. 2017), degenerated retinal function (Sekirnjak et al. 2009), and retinal activity evoked by a photovoltaic subretinal prosthesis (Mathieson et al. 2012; Wang et al. 2012; Lorach et al. 2015a; Goetz et al. 2015; Mandel et al. 2013; Boinagrov et al. 2014; Lorach et al. 2015b). The approach yields explicit quantitative predictions of visual perception of images that are consistent with two known aspects of prosthetic vision. First, the predicted image obtained with a retinal prosthesis is severely degraded compared to the image obtained in normal vision over the same field of view, both in healthy and degenerated retina simulations. This finding is consistent with the limited resolution and lack of cell type specificity of existing prostheses. Second, re-optimizing reconstruction for prosthesis stimulation substantially improves the reconstructed image. This indicates that the potential for learning by patients to improve prosthesis function is high. We comment on potential enhancements to make the approach more realistic, on the use of this framework for design of future prosthesis technology, and on the potential for image processing to optimize the function of existing devices.

## Methods

### Overview

A computational framework was developed to infer a visual percept based on RGC activity, both for healthy light activation, and for electrical activation using a subretinal prosthesis. The <u>ISETBio</u> (Image Systems Engineering Toolbox – Biology) software was used to model transformation of a visual stimulus into RGC spikes in the healthy retina (Fig. 1). Additions to software were used to simulate electrical activation of bipolar cells via a retinal prosthesis, in turn producing RGC spikes. A linear transformation of simulated RGC responses was then used to infer the perceived image, over samples of a large image data set. The linear transformation was selected to optimize reconstruction for either the healthy or prosthesis condition.

**Figure 1.**
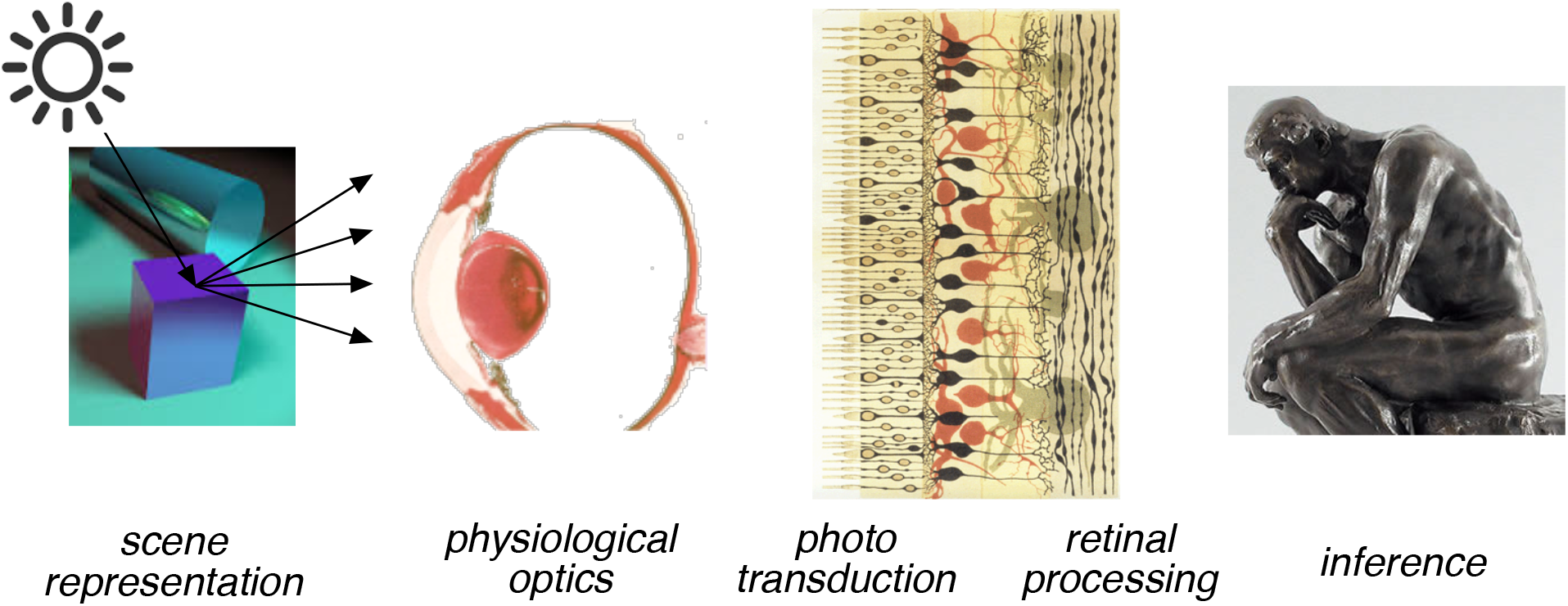
The ISETBio computational pipeline. Stimuli represented by spectral radiance over space are transformed to retinal irradiance, cone photopigment isomerizations, photocurrent, bipolar responses, and finally RGC spikes, which are then used for linear reconstruction or classification (Jiang et al. 2017). Reprinted with permission of IS&T: The Society for Imaging Science and Technology, sole copyright owners of the *Electronic Imaging, Human Vision and Electronic Imaging 2017.*

### Healthy Retina Simulation

Simulations of signals in healthy retinas were produced using the ISETBIO software, a Matlab toolbox that allows one to estimate the biological representation of a visual stimulus at different stages of visual encoding (Cottaris et al. 2018). The ISETBio simulation begins with a representation of image radiance. This is transformed into retinal irradiance via the simulated cornea, lens, and inert pigments. Irradiance is then transformed into a spatial array of photon absorptions and photocurrent for each cone in the photoreceptor mosaic. Cone photocurrents are then linearly transformed into bipolar responses, with appropriate cone type connectivity. Finally, bipolar responses are transformed into RGC responses via a generalized linear model that produces simulated spikes (Pillow et al. 2008). This package simulates many aspects of healthy retinal responses to arbitrary visual stimuli, although it does not yet explicitly include the role of horizontal and amacrine cells or diverse nonlinear aspects of retinal processing. Below, the full spatio-temporal model is described. Later, the more limited use of the model in simulating static image presentations, RGC responses, and reconstruction is presented.

The stimulus image was modeled as an array of MxN pixels on a RGB computer display, and summarized by an MxNx3 matrix. The spectral power density of the display primaries was used to calculate a larger array (MxNxW) representing the display radiance, where each value of W indexes the radiance at a particular wavelength. The full spectral image was then passed through a model of the human optics based on mean wavefront aberrations measured on-axis (near fovea) as well as lens and macular pigment densities. The retinal spectral irradiances were used to compute the Poisson distributed photopigment isomerization rates for the L, M and S cones in the simulated cone mosaic. Cone isomerizations were computed using photopigment parameters for human foveal vision, which when combined with the assumed lens and macular pigment densities reproduce the CIE foveal cone fundamentals (CIE 2006). A linear temporal filter was applied to the cone isomerizations to generate simulated cone photocurrents over time (Angueyra and Rieke 2013).

Four types of bipolar cells were modeled, ON diffuse, OFF diffuse, ON midget, and OFF midget. Each type pooled inputs linearly over a local collection of cones. The bipolar temporal response was assumed to be equal to that of the RGC type to which it provided input (see below). RGC temporal responses were obtained from a standard data set (Pillow et al. 2008). Thus, the bipolar temporal response was obtained by convolving the cone response in time with the additional temporal filter required to transform the cone response into the temporal response of the appropriate RGC type. The connectivity of the three cones types to the four types of bipolars was modeled as follows: diffuse and ON midget bipolar cells received input from only L and M cones, while OFF midget bipolar also received input from S cones with a magnitude 25% that of L and M cones (Dacey 2000; Field et al. 2010).

The RGC response was simulated with a generalized linear model based on bipolar cell input (Pillow et al. 2008). The four bipolar types above provided the sole input to ON parasol, OFF parasol, ON midget and OFF midget RGCs, respectively. The input to the RGC was computed as a local sum of bipolar cell responses, chosen so that the stimulus-referred spatial properties of the RGC fitted a published data set (Pillow et al. 2008). Because the bipolar temporal response was already chosen to equal that of the RGC, the transformation from bipolar to RGC did not involve further temporal filtering. The input to the RGC from bipolars was then followed by exponentiation and Poisson spiking with a post-spike filter, obtained from fits to experimental data (Pillow et al. 2008).

The simulated patch of retina was a square region 1.7° (512 μm) on a side, centered at 1.8° eccentricity. The simulated patch had a uniform density of cones, bipolars, and RGCs appropriate for 1.8° eccentricity, and did not account for the changing density of these cells over the field of view. The cone mosaic over the region was a 256×256 square lattice with a pitch of 2 μm. The mosaics of each bipolar cell type had the same dimensions as the cone lattice. Each bipolar cell integrated over cone inputs using a difference of Gaussians spatial weighting function with center peak amplitude of 1, surround peak amplitude of 0.01, center SD of 4 μm, 4 μm, 2 μm and 2 μm for the four types of bipolar cells respectively, and surround SD equal to 9 times the center SD (Dacey 2000). The RGCs in the simulation consisted of approximately hexagonal lattices of 28×32, 31×35, 55×63, and 61×70 cells for the four RGC types, respectively, yielding a total of 9,716 RGCs. The four RGC types received inputs from nearby bipolars, weighted according to a difference of Gaussians function with center peak amplitude of 1, surround peak amplitude of 0.375, 0.375, 0.5 and 0.5 for the four RGC types, center SD of 10 μm, 8 μm, 4 μm, and 3.5 μm, and surround SD equal to 1.15 times the center SD. These parameters were selected to approximately match experimental data (Pillow et al. 2008; Croner and Kaplan 1995). Poisson spiking by RGCs was the only source of noise in the simulation.

### Degenerated Retina Simulation

Conditions like retinitis pigmentosa and macular degeneration cause structural changes in retinal circuitry. In order to model their effects on signals in the bipolar and retinal ganglion cell populations, three of the observed physiological effects of degeneration were included in the model.

First, in the healthy retina, bipolar cells of a particular subtype form connections with only one subtype of retinal ganglion cell, but in a degenerated retina they sometimes form connections with other bipolar cells, including those of different types (Marc et al. 2007, Jones et al. 2012, Wong et al. 2016). This was implemented by adding stochastic connections between nearby bipolar cells of all types with a 5% probability.

Second, changes in spontaneous activity of retinal ganglion cells have also been observed in the degenerated retina, notably a significant increase in the firing rate of OFF cells (Sekirnjak et al. 2009). The model captures this with a 20% larger constant in the Poisson process that simulates spontaneous spiking.

Third, the fraction of ganglion cells that survive and respond to prosthetic stimulation also decreases significantly during degeneration, across cell types. This was implemented with random dropout of 30% of ganglion cells in the reconstruction process, reflecting observations from moderate retinitis pigmentosa (Santos et al., 1997).

### Subretinal Prosthesis Simulation

To model the encoding of the visual signal by a subretinal prosthesis, a simulation of the electrical stimulation of bipolar cells was introduced into the ISETBio pipeline. The simulation was modeled after a subretinal prosthesis, in which a visual stimulus captured by a camera mounted on goggles is transformed into pulses of infrared light that irradiate a grid of photovoltaic pixels implanted in the sub-retinal space near the bipolar cells (Mathieson et al. 2012). In the simulation, an 8×8 hexagonal array of photodiodes with 70 μm pitch covered the field of view. The voltage of each photodiode at each time step was assumed to be proportional to the irradiance of the visual stimulus integrated over the photodiode area during that time step. The electrodes produced responses in nearby bipolar cells in proportion to this voltage. The spatial profile of bipolar activation was the profile of the grid of electrode center points, each scaled according to stimulation voltage, convolved with a Gaussian function with a SD of 35 μm. Each bipolar cell was activated in proportion to the value of this activation profile at its location. The constant of proportionality was set to approximately mimic the distribution of RGC responses that occurs with natural scene stimulation (data not shown). With this constant, the stimulus contrast required for reliable detection of a full-field step of light (see below) was 3.1%. The temporal response of the bipolar was given by the activation value convolved with the bipolar temporal impulse response. This implementation did not capture nonlinear effects such as saturation of the photovoltaic pixels at high input irradiances (Mathieson et al. 2012).

### Static Simulation

Although the calculations above yield simulations of RGC activity over time, most of the analysis was performed for static images and static RGC responses, for simplicity. In both the healthy retina and prosthesis simulation, images were presented for 0.4 s, and the responses of RGCs were accumulated over the time window 0.320-0.328 s after the stimulus presentation. This produced a vector of spike counts across the RGC population, in response to each static image.

### Optimal Linear Reconstruction

Linear reconstruction was used to estimate the visual information available to a patient based on RGC activation. To perform linear reconstruction, a set of spatial filters were first computed for the RGC population. Each spike emitted by a RGC during the response caused a copy of the spatial filter to be placed in the reconstructed image. The final reconstructed image was the sum of the filters contributed by all spikes from all RGCs. The spatial filters for the collection of RGCs were computed under the constraint that, when applied to RGC responses in either healthy or prosthesis simulation, they produced reconstructed images with least squared deviation of pixel intensity from the original stimulus. The identification of optimal filters was performed on 80% of the images in the data set. The performance of reconstruction was then assessed on the remaining 20% of images. In total, 288,000 image patches were used, each 100×100 pixels, sampled from the Imagenet database (Deng et al. 2009). Note that the optimal filters depend on both the statistics of the stimulus and correlations in the population response. For natural scene stimuli, the spatial reconstruction filter and the spatial receptive field for a particular cell do not necessarily match, although they are usually similar in shape and located in the same region of the visual field (see below, Fig. 8).

Reconstruction filters were calculated using linear regression (Warland and Meister 1997). Let s denote the stimulus image as a row vector, let r denote a row vector indicating the number of spikes from each RGC with an offset term of 1 as its first entry, and let W denote the reconstruction matrix, in which each row contains the reconstruction filter for a particular RGC. The reconstruction srec of the stimulus s is then given by:

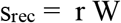

To obtain a matrix W that reconstructs as accurately as possible for a collection of images, let S be a matrix containing an image in each row and let R be the matrix containing the corresponding responses from all cells in each row, and let Srec be a matrix containing the reconstructed images in each row. The objective is to minimize the squared difference between reconstructed and original images,

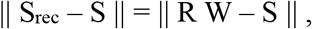

where ||•|| represents the sum of squared entries. This is a standard regression problem with the solution (Menke 2012):

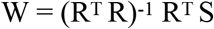

The reconstruction filters in the columns of W account for correlations in the firing of nearby RGCs, which causes the filters to differ from the individual receptive fields obtained from spike-triggered averaging. The performance of reconstruction using the matrix W was then assessed on a set of held out stimuli and responses. As expected, reconstruction performance was limited by the amount of data used to identify the entries of W. To obtain robust estimates of W, regularization was performed by limiting the rank of W (Chambers 1977). Specifically, if R is given by its singular value decomposition,

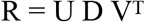

Then, from the expression for W above, we have:

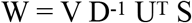

Thus, an approximation to W with rank K is given by:

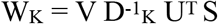

Here D^-1^_k_ denotes zeroing all but the first K diagonal entries of D^-1^. For the healthy retina reconstruction, retaining the top 50% of singular values (K=4904) produced optimal performance. For the prosthesis reconstruction, retaining the top 5% of singular values (K=491) produced optimal performance. These values were used in all analyses above.

### Linear Classification for Comparison to Clinical Tests

In order to simulate measurements of perceptual thresholds, such as those performed on patients with retinal implants (Stingl et al. 2013), a linear classification approach was used (Farrell et al. 2014). Two sets of stimuli were processed by the prosthesis simulation, producing reconstructed images, and a linear SVM classifier was trained to discriminate pairs of reconstructed images, one from each set. Classification performance was then assessed on held out data as a function of a particular stimulus variable, e.g. contrast or spatial frequency. Discrimination thresholds were defined as the value at which the classifier achieved 82% accuracy.

## Results

### Simulation of retinal light response and prosthesis activation

To quantify normal visual system function, as a benchmark for artificial vision with a prosthesis, a simulation of the normal (healthy) light activation of retinal ganglion cells (RGCs) by visual stimuli was produced using the publicly-available <u>ISETBio</u> software (see Methods). Briefly, each RGC was modeled using a generalized linear model (Pillow et al. 2008), with center-surround receptive field organization (Kuffler 1953). Simple cascade models have been used to capture the responses of populations of RGCs in previous studies (Chichilnisky 2001, Chichilnisky and Kalmar 2002, Field and Chichilnisky 2007, Pillow et al. 2008). The simulation was built on modeled inputs from photoreceptors transmitted through bipolar cells in the retinal network. The simulation focused on the four numerically dominant RGC types in macaque and human retinas: ON parasol, OFF parasol, ON midget, and OFF midget (Dacey 1999). Each of the four cell types forms a mosaic uniformly covering the visual scene (Fig. 2, top). The spatial, temporal, and nonlinear contrast-response properties of each RGC type (Fig. 2) were taken from experimental findings. Presentation of a static natural visual image (Fig. 3, left) to the simulated retina caused a spatial pattern of activation in the four RGC types (Fig. 3, second and third columns). Although the simulation produced RGC spikes over time, for simplicity most of what follows focuses on flashed static images reconstructed from total RGC spike counts. Dynamic stimuli and responses are considered briefly at the end of Results.

**Figure 2.**
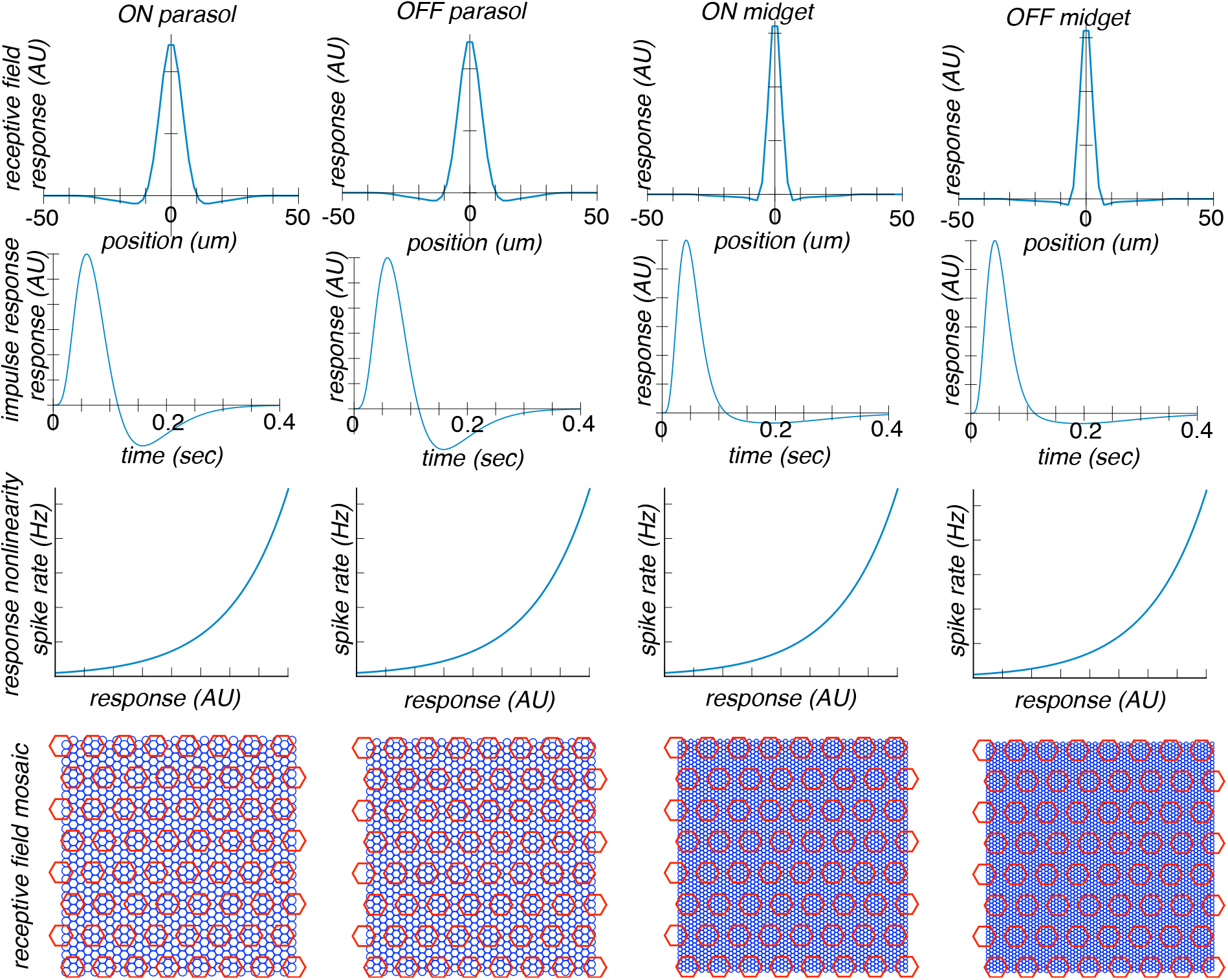
Simulation of RGC responses. Each RGC is modeled as a linear-nonlinear cascade (see Chichilnisky 2001, Chichilnisky and Kalmar 2002), with an intermediate representation of photoreceptor and bipolar cells (see Methods). Top row: cross-sectional spatial profiles of the spatial receptive fields of each RGC type, with midget cell RF extents smaller than parasol, and OFF slightly smaller than ON. Second row: the temporal impulse response of each RGC type; a typical biphasic response, with different details in midget and parasol RGCs. Third row: the response nonlinearity of each RGC type is an accelerating half-wave rectifying function. Bottom row: the collection of RFs of each RGC type (blue circles) uniformly tile the region of retina, with different densities, and the simulated implant electrodes (red circles) form a much coarser sampling of space.

To simulate the retinal signal that would be produced by a prosthesis, a model was developed of the activation of the retinal network by the electrodes of a wireless subretinal photovoltaic device (Loudin et al. 2007). This technology has been tested extensively in isolated retina as well as *in vivo* animal studies, and is now being tested in clinical trials (Goetz et al. 2015; Mandel et al. 2013; Boinagrov et al. 2014; Lorach et al. 2015b). In the simulation, the presentation of a static visual image to the prosthesis produced spatial patterns of current from the electrodes. This current then caused depolarization of the various bipolar cell types that provide input to the RGCs, resulting in spiking activity in the RGC populations (Fig. 4B, C, D, E). The pattern of activation induced by the prosthesis was quite different from the activation of RGCs in the healthy retina simulation (Fig. 3B, C, D, E). A salient difference is that the normal opposite activation of ON and OFF populations by light and dark portions of the visual scene was not reproduced, because light-evoked current in the prosthesis activates ON and OFF bipolar cells indiscriminately and with the same polarity. This lack of cell type specificity is a known limitation of existing prosthesis technology (Goetz et al. 2015; Goetz and Palanker 2016; Jepson et al. 2013, 2014).

**Figure 3.**
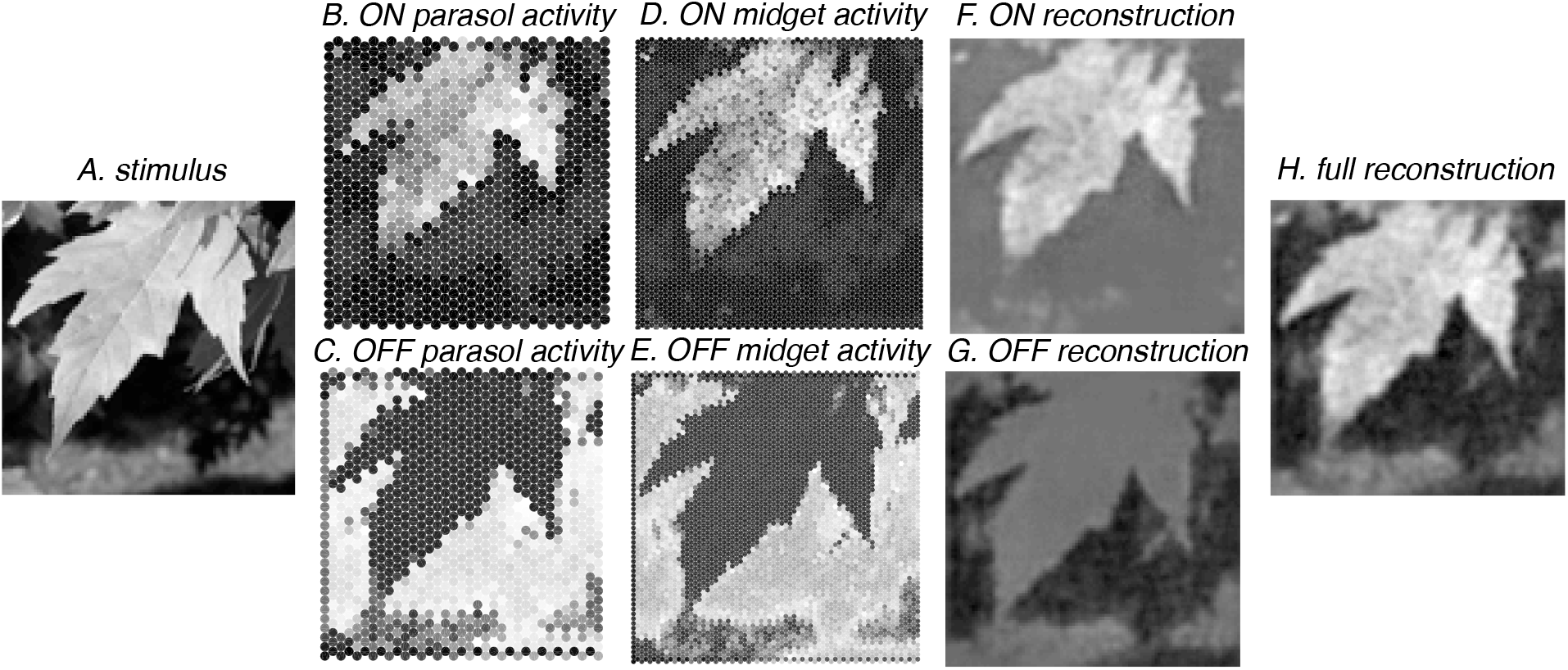
Simulation of visual stimulus, response, and linear reconstruction in healthy retina. **A**: a sample visual stimulus. **B, C, D, E**: simulated responses of the four major RGC types over space to the flashed stimulus, with light colors indicating higher activity. **F, G**: linear reconstruction of the visual input by ON cells and OFF cells, generated by applying the spike responses shown in the second and third columns to the reconstruction filters trained on a large collection of responses to natural images. **H**: linear reconstruction of the original stimulus from all four RGC types combined.

Patients receive prosthetic implants because of a loss of photoreceptor cells, which results in blindness. However, the retina is also known to undergo substantial remodeling (Marc et al. 2007, Jones et al. 2012) and death of other neurons (Santos et al. 1997). A degenerated retina model was therefore developed that differs in three ways from the healthy model: (1) the OFF-type RGCs had a 20% elevated spontaneous firing rate, (2) the bipolar cell mosaics had random connections with bipolar mosaics of other cell types with probability 5%, and (3) 30% of RGCs did not fire at all (see Methods). These effects, most notably the cells that do not fire, are evident in the activation patterns of the RGCs (Figs. 5B, C, D, E). Note that although these values may approximate the effects of moderate retinitis pigmentosa, clinical presentation varies widely, and some patients have more extensive damage that leads to the loss of up to 70% of RGCs (Santos et al. 1997).

### Linear reconstruction

To assess how the different patterns of RGC activity with normal and prosthesis stimulation may influence visual perception, a linear reconstruction of the visual stimulus was generated based on activation of RGCs. The reconstructed images provide an estimate of the stimulus information that is available to the patient. Specifically, in the simulation, the pattern of RGC activation is linearly transformed into a simulated “perceived” image that deviates as little as possible from the original. This deviation was measured as the mean squared error across an ensemble of natural images (Warland 1997, Rieke 1997, Nirenberg and Pandarinath 2012). These key technical assumptions (linear reconstruction, squared error, natural images) are considered further in Discussion. Based on these assumptions, image reconstruction is a regression problem, in which an optimal linear transformation is calculated that transforms RGC activation to the estimated incident image. Regression was performed using 230,400 natural images and responses to them (see Methods); the transformation was then evaluated on 57,600 held-out images. Conceptually, this linear transform amounts to assigning a fixed spatial filter to each RGC (Fig. 8, left), such that each spike from the RGC contributes that filter to the reconstructed image, and contributions from all spikes and RGCs are summed. The filter for each cell can be understood as representing the visual information conveyed by a spike in that cell.

In the case of natural activation of RGCs, linear reconstruction produced an image that resembles the original image (Fig. 3H), with a resolution limited by the density of RGCs. Because the reconstruction is linear, the contributions of the different RGC types can be examined separately. For example, ON cells provided similarly detailed contributions to the reconstruction as OFF cells (Fig. 3F, G).

In the case of prosthetic activation of RGCs, the reconstruction procedure produced a degraded image, in which few features of the original image are easily recognizable (Fig. 4H). This degradation is not surprising given that the ON and OFF bipolar cells, and thus the ON and OFF RGCs, are activated similarly by the simulated prosthesis (Fig. 4B, C, D, E). As a consequence, because spikes from ON and OFF RGCs (Fig. 3B, C, D, E) are treated oppositely in reconstruction optimized for the normal retina (Fig. 3F, G), their contributions to the full reconstructed image would often be expected to cancel. This interpretation was confirmed by examination of the ON and OFF reconstructions separately: the two groups of cells each produced more accurate reconstructions (ignoring the sign of the reconstructed contrast) individually (Fig. 4F, G) than when combined. Averaged over test images, the root mean square (RMS) error for reconstruction with all cells was 0.09 in the normal retina simulation, and 0.22 for prosthesis stimulation. Note that this model does not include the possible effects of retinal degeneration, which will be addressed below. As a comparison, the RMS error obtained with temporally shuffled spikes bearing no relationship to the visual stimulus was 0.24. Thus, a computation that assumes ON and OFF RGCs are equally stimulated, and that their signals are used to reconstruct the visual scene as they would be for healthy retina, predicts that the perceived visual image with a prosthesis is severely degraded.

**Figure 4.**
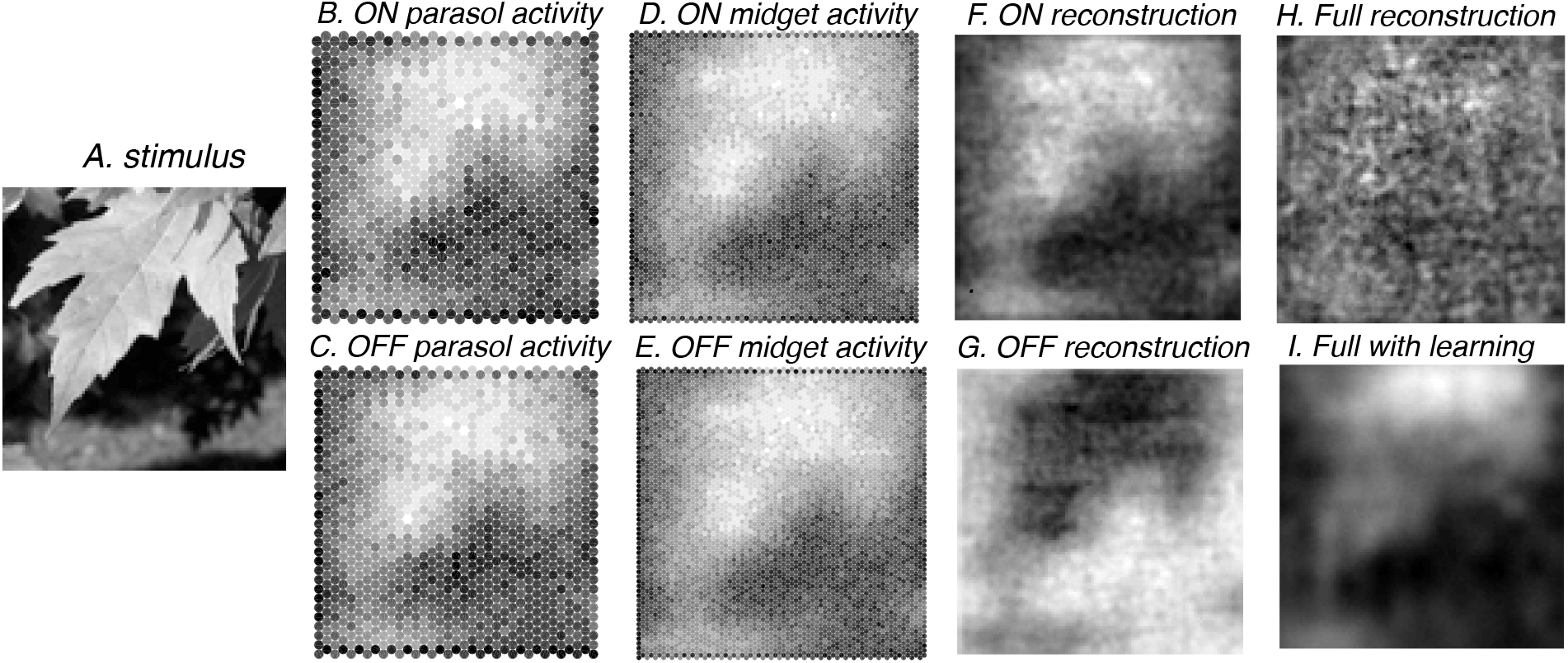
Simulation of stimulus, response, and linear reconstruction with retinal prosthesis. **A**: visual stimulus. **B, C, D, E**: simulated responses of four major RGC types to stimulation with the prosthesis. Note the similar patterns of activation of ON and OFF cells by the prosthesis. **F, G**: linear reconstruction of the stimulus, using the reconstruction filters obtained from the healthy retina simulation, separately using ON and OFF cells. Note the reverse polarity of the reconstruction for OFF cells. **H**: reconstruction with all four RGC types combined normally, causing cancellation of ON and OFF signals. **I**: reconstruction with all four RGC types combined, using reconstruction filters obtained with prosthesis stimulation, mimicking optimal learning by the patient.

Under these assumptions, factors that might otherwise be expected to improve prosthesis function, namely the density of electrodes and magnitude of current spread, had little impact on reconstruction performance. For example, reducing electrode spacing and current spread by a factor of two produced a decrease in RMS error of only 4.5% (Fig. 6E, regression line). Thus, the effects of nonspecific activation can in principle be so severe that incremental technical advances in prosthesis design would be of little value.

The naive prediction of severely degraded artificial vision above appears inconsistent with patient reports from existing prosthesis technologies, which indicate that some degree of spatial vision is in fact possible (see Goetz and Palanker 2016). One possibility, suggested by the dominance of “bright” (rather than “dark”) phosphenes reported by patients (Stingl et al. 2013), as well as by hypothesized asymmetries in electrical activation of ON vs. OFF cells (Werginz et al. 2015; Cottaris and Elfar 2005), is that the contribution of ON cells to artificial vision is stronger than that of OFF cells, reducing the problematic cancellation of visual signals described above. To simulate this hypothesis in an extreme form, reconstruction was performed using only ON cells (Fig. 4F), with no contribution from OFF cells. In this simulation, the average RMS reconstruction error across images was 0.14 (Fig. 6F), a value less accurate than in the healthy condition (0.09), but more accurate than was obtained using both ON and OFF RGCs (0.22). Note, however, that in this condition the additional improvement in performance provided by higher resolution stimulation was still only 8% (Fig. 6F). In sum, the artificial vision provided by retinal prostheses in the clinic could benefit substantially from a dominance of specific cell types, though large improvements in spatial resolution may still be difficult to achieve.

### Degenerated Retina

How would changes in retinal circuitry that occur during degeneration affect the ability to accurately reconstruct the image from prosthesis stimulation? This was tested by simulating the degeneration process as described above.

The effect of simulated degeneration can be observed in the patterns of RGC activation (Fig. 5B, C, D, E), which show a marked difference from the patterns for the healthy retina under prosthesis stimulation (Fig. 4B, C, D, E). The similarity between the activation patterns and the stimulus is much less apparent with simulated degeneration. As in the healthy retina condition, reconstruction with the full response was inaccurate (Fig. 5H), consistent with non-specific activation of ON and OFF cells. Reconstruction using only ON cells in the degenerated retina was also inaccurate (Fig. 5F) with an RMS error of 0.20 (Fig. 7F), substantially higher than the value in the healthy retina simulation (0.14). In summary, the effects of simulated degeneration are severe, causing substantial additional degradation of reconstruction beyond the degradation produced by nonspecific activation with the prosthesis.

**Figure 5.**
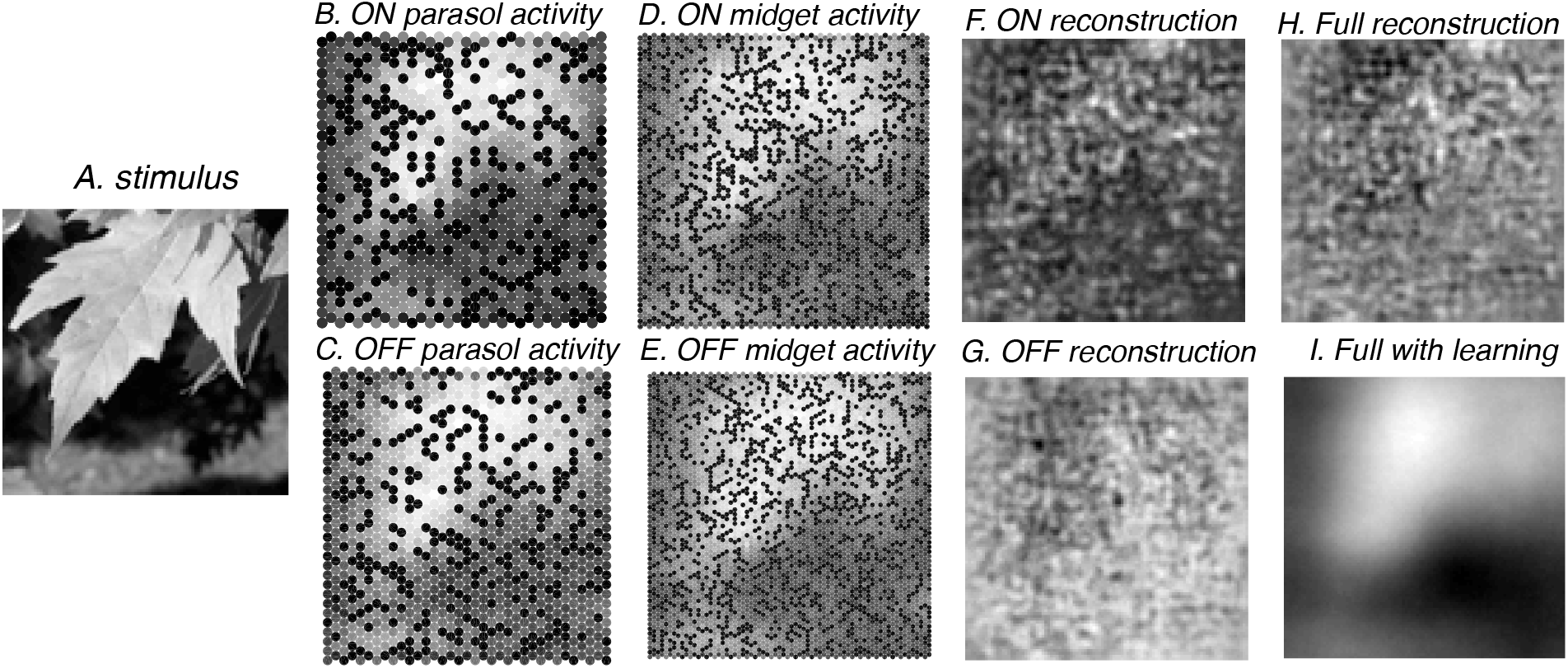
Simulation of stimulus, response, and linear reconstruction with prosthesis stimulation of degenerated retina. **A**: visual stimulus. **B, C, D, E**: simulated responses of four RGC types to prosthesis stimulation in degenerated retina model. **F, G**: linear reconstruction using filters from the degenerated retina simulation. 30% of RGCs do not fire. H: reconstruction with all four types, again causing cancellation between ON and OFF signals. I: reconstruction with all types combined, using filters from the degenerated simulation with learning.

In the case of degeneration, two-fold reduction in electrode spacing and current spread improved reconstruction by 7% (Fig. 7F), close to the improvement seen in the healthy retina simulation (8%, Fig. 6F), suggesting that advances in prosthesis technology could produce similar clinical benefits for the degenerated retina.

**Figure 6.**
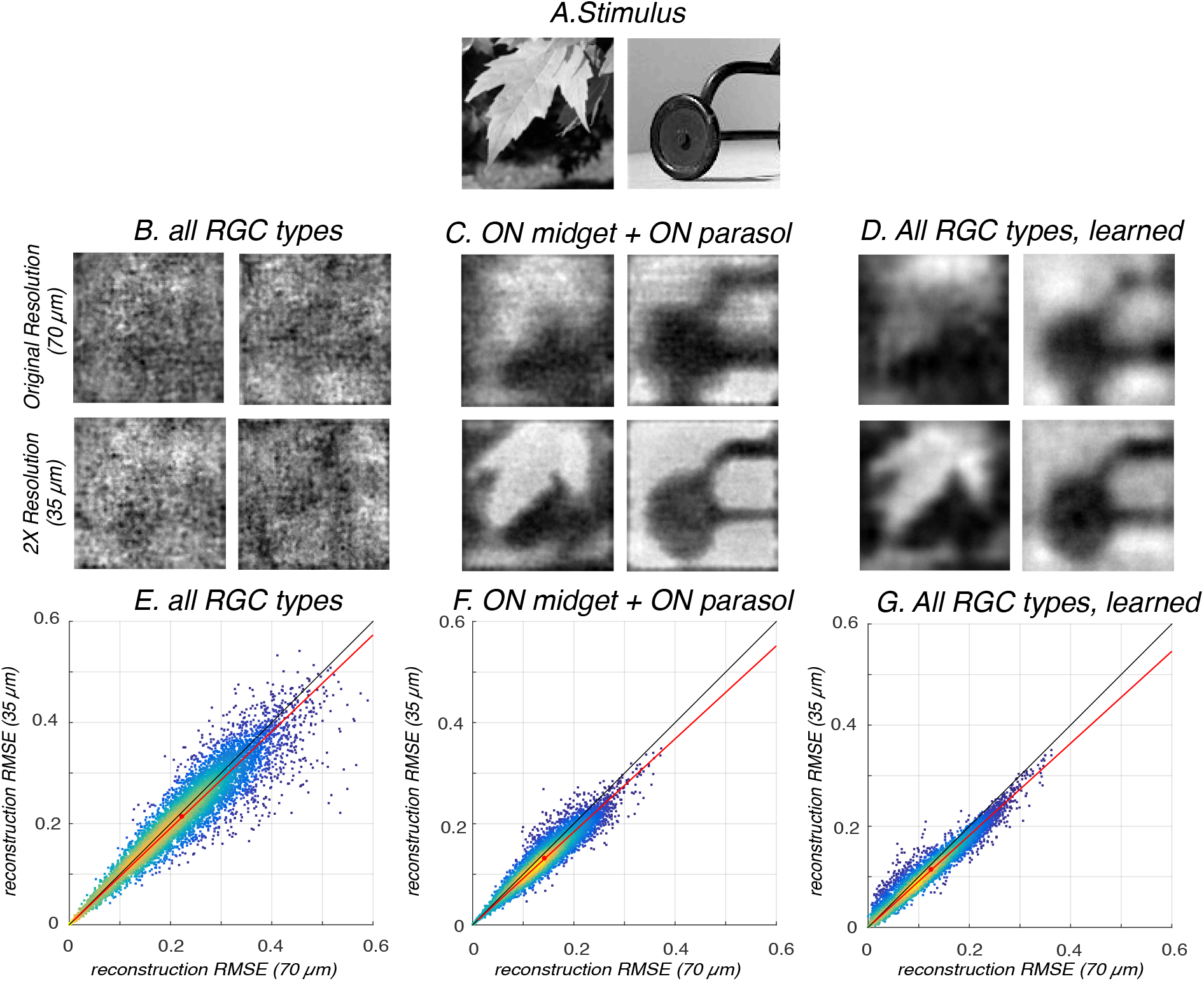
Reconstruction performance with denser electrodes and lower current spread. **A**. Visual stimuli. **B, C, D**: Example reconstructions. **E, F, G**: the root mean square error (RMSE) for reconstruction of 57,600 tested images, using the original electrode spacing and current spread (70 μm, 35 μm) vs. values obtained with both reduced two-fold (35 μm, 17.5 μm). **E**: reconstruction error using all four RGC types. Red line shows linear regression to the data; regression slope of 0.955; red circle shows mean value of 0.22 for 70 μm and 0.21 for 35 μm. **F**: reconstruction error using only ON cells; regression slope 0.92 for the prosthesis; red circle shows mean value of 0.14 for 70 μm and 0.13 for 35 μm. **G**: reconstruction error using all four RGC types, with learning; regression slope 0.91; red circle shows mean value of 0.12 for 70 μm and 0.11 for 35 μm.

### Learning

Could artificial vision improve if the brain learns to interpret the unnatural patterns of RGC activation that occur with a prosthesis? To answer this question, optimal reconstruction filters were re-calculated using RGC activity elicited by the prosthesis, instead of natural RGC activity, in both the healthy and degenerated retina models. This simulation reveals how much improvement in stimulus representation can be achieved if the reconstruction is optimized to the response pattern created by the prosthesis (i.e. optimal “learning”). As expected, the reconstruction filters associated with RGCs were very different in the learning simulation (Fig. 8), because the electrically stimulated RGC activity used for learning was very different from the healthy RGC activity. Notably, reconstruction filters exhibited asymmetric shapes that varied between cells, due to misalignment between electrodes and receptive fields of stimulated cells. Also, OFF cells often exhibited filters with dominant ON polarity, consistent with the polarity of prosthesis stimulation. In other words, learning produced a reconstruction that most effectively exploited the signals originating from different electrodes in the prosthesis, which vary because of the different cells and cell types stimulated by each electrode. This learning can be interpreted as the patient’s central visual system optimally utilizing the unnatural signals produced in RGCs by the prosthesis.

With learned filters, reconstruction of images using all RGC types was substantially improved (Fig. 4I; Fig. 6G). The RMS error of reconstruction across test images was 0.12 (Fig. 6G), closer to the value achieved in natural vision (0.09) than was observed with only ON reconstruction (0.14), as expected. In this condition, a two-fold reduction in electrode spacing and current spread reduced RMS error by 9% (Fig. 6G), similar to the improvement seen in the only ON reconstruction (8%).

For the degenerated retina, learned filters also allowed for a substantial improvement in reconstruction with all RGC types, producing a RMS error of 0.13 (Fig. 7G), much lower than the original full reconstruction error (0.24). In this condition, there was a 10% improvement with two-fold reduction in electrode spacing and current spread (Fig. 7G).

**Figure 7.**
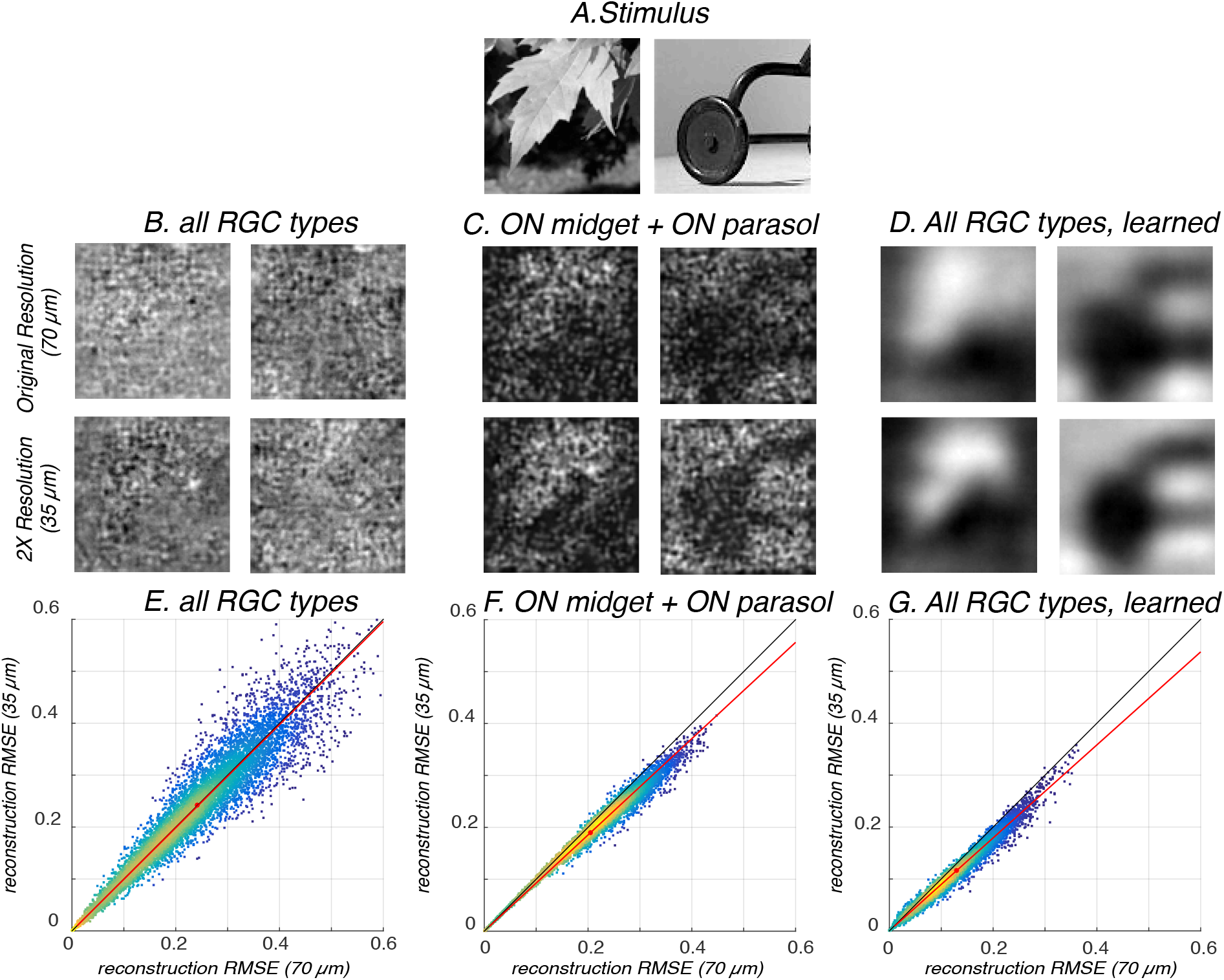
Reconstruction performance with denser electrodes and lower current spread for the degenerated retina. **A**. Visual stimuli. **B, C, D**: Example reconstructions. **E, F, G**: the root mean square error (RMSE) for reconstruction of 57,600 tested images, using the original electrode spacing and current spread (70 μm, 35 μm) vs. values obtained with both reduced two-fold (35 μm, 17.5 μm). **E**: reconstruction error using all four RGC types. Red line shows linear regression to the data; regression slope of 0.99; red circle shows mean value of 0.24 for 70 μm and 0.24 for 35 μm. **F**: reconstruction error using only ON cells; regression slope 0.93 for the prosthesis; red circle shows mean value of 0.20 for 70 μm and 0.19 for 35 μm. **G**: reconstruction error using all four RGC types, with learning; regression slope 0.90; red circle shows mean value of 0.13 for 70 μm and 0.12 for 35 μm.

In summary, the simulation framework predicts that the *potential* for patients to learn the unnatural neural code and improve prosthesis function is significant, including in the degenerate retina. Such learning would also enhance the benefits from technical improvements in the device.

### Simulated behavioral tests

A potential application of the reconstruction framework to functional assessment in the clinic is the prediction of visual performance on standardized tasks, including discrimination of Landolt C patterns and grating orientation as a function of spatial frequency at a set contrast (100% for the Landolt C and 4% for the gratings). To predict performance in these tasks, visual images were presented to the simulation, and reconstruction was performed as above. Then, the reconstructed images were used to classify the stimulus, using a linear classifier trained on reconstructions obtained from many presentations of known stimuli (see Methods). For example, in the Landolt C task, stimuli containing a gap were discriminated from stimuli not containing a g^a^p.

At a set contrast, the discrimination accuracy progressively increased with gap size (Landolt C) and decreased with spatial frequency (gratings), as expected (Fig. 9). However, the different simulations showed differences for the two stimuli as well. For example, the ON-only performance was similar to “learned” performance for in the Landolt C experiment, while it was generally lower and depended less strongly on spatial frequency in the gratings experiment. Thus, simulations reveal that certain visual stimuli may be more effective than others in inferring the mechanisms mediating successful prosthetic vision in the clinic.

**Figure 8.**
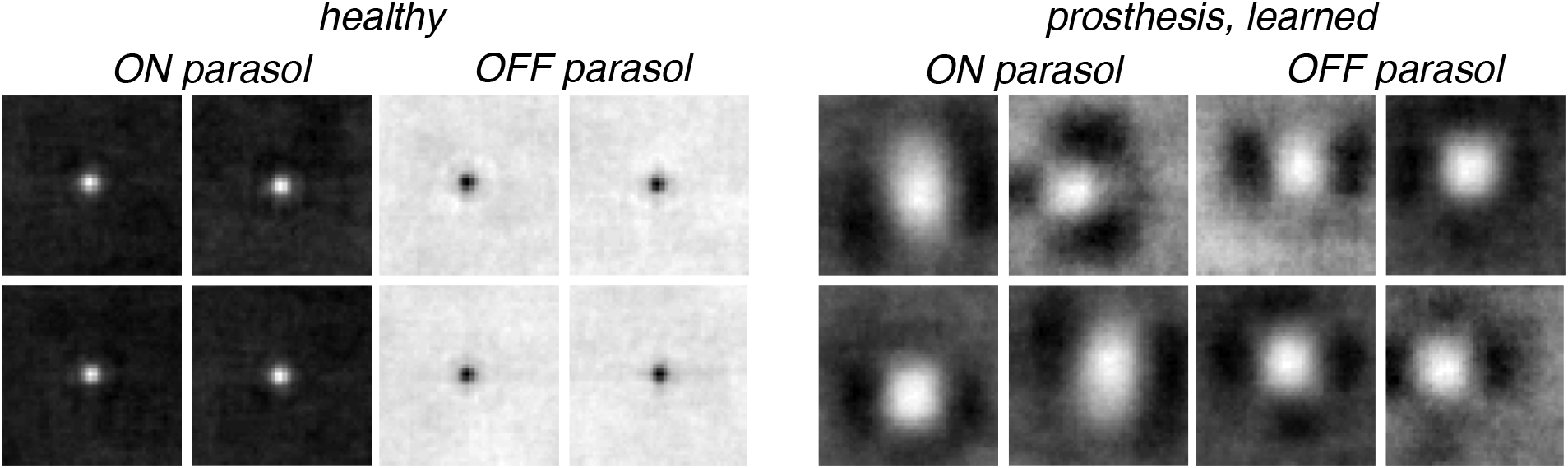
Reconstruction filters in healthy and prosthesis simulations. Left: sample reconstruction filters are shown for ON and OFF parasol cells (four each) obtained from the healthy retina simulation. The filters roughly resemble the spatial receptive fields of the cells (not shown), and their center polarity corresponds to the polarity of light response. Right: sample spatial reconstruction filters for the same cells obtained from the prosthesis simulation. In this case, reconstructions filters are asymmetric and variable, and largely reflect the ON polarity of prosthesis stimulation for both cell types. Each image is centered on the corresponding cell location.

**Figure 9.**
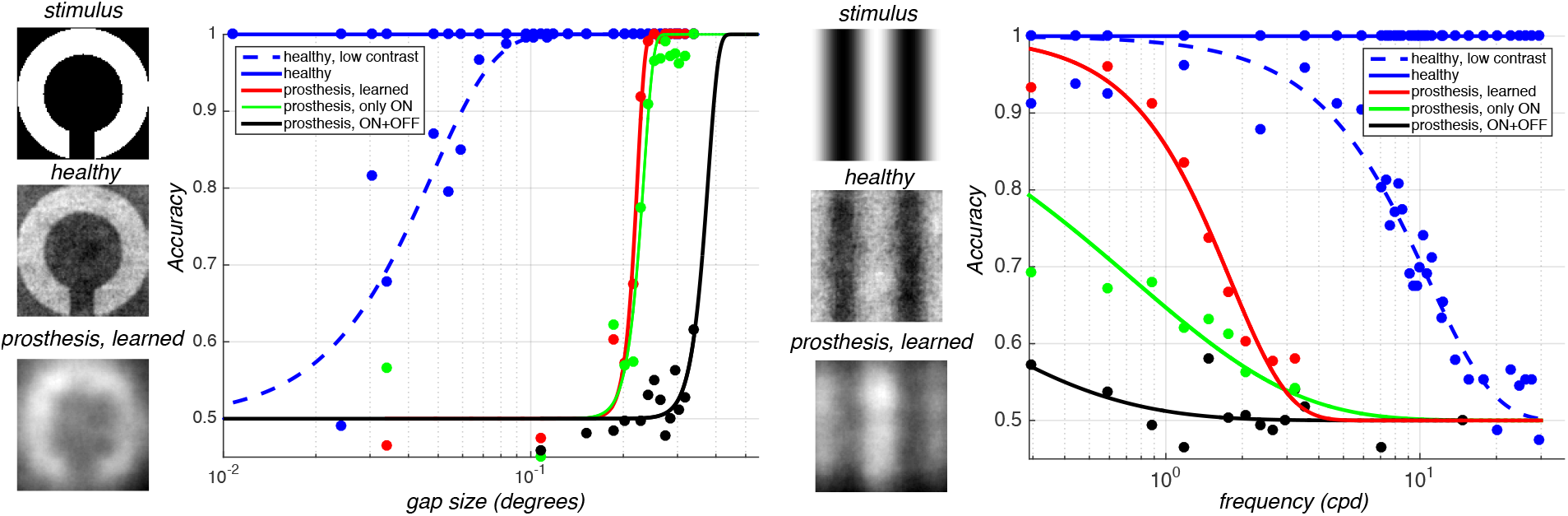
Landolt C and grating orientation discrimination, using healthy and prosthesis with degenerated retina simulations. Left: Landolt C stimuli, and similar stimuli lacking a gap, were presented to the simulation, producing reconstructions for healthy retina, prosthesis-stimulated retina with learning, prosthesis-stimulated retina with no learning for only ON cell responses, and prosthesis-stimulated retina with no learning for ON and OFF cell responses. Accuracy of linear classification of stimuli with and without gaps decreased as a function of the visual angle of the Landolt C gap at a contrast of 100%. Classification was most accurate for the healthy retina simulation, an order of magnitude less accurate for the prosthesis with learning or no learning and only ON responses, and another factor of two less accurate for the prosthesis with no learning for ON and OFF responses. Note that for the healthy condition, accuracy was perfect for all gap sizes, so a low contrast Landolt C stimulus at 4% was used to observe the psychometric function (dashed line). Right: Grating stimuli, horizontally or vertically oriented, were presented to the simulation and discriminated. Similar large discrepancies in the different conditions were observed. Note the unrealistically high accuracy at low contrast, reflecting a highly-trained linear classifier well aligned with the visual stimulus (see Methods). Accuracy generally decreased as a function of spatial frequency for a 4% contrast gratings stimulus, with the ordering of accuracy of each condition resembling that in the Landolt C simulations. For the healthy condition, accuracy was perfect for all spatial frequencies at 4% contrast, so a relatively lower contrast stimulus of 0.08% was used to observe a psychometric curve for the healthy response (dashed line).

Note that healthy orientation discrimination performance was unrealistically high compared to normal human performance: patients have been tested for gratings at a contrast of 100%, and many fail to discriminate between stimuli at any spatial frequency (Stingl et al. 2015, Stingl et al. 2013). This may be in part because the classification simulation has unrealistic prior “knowledge” of the stimulus pattern and location, and because the retinal simulation captures only some of the biological noise introduced in the retina and none of the noise introduced downstream of the retina. Thus, a meaningful quantitative prediction of absolute behavioral performance would require a more realistic simulation of visual discrimination, including consideration of the size of the stimulus, the task, the nature of the classifier applied to the simulated output, and assumptions about noise in the retina and the central visual system. The present results, instead, mainly point out relative differences in expected performance across different stimuli.

### Dynamic reconstruction

The above simulations were all performed using only static spatial images and integrated RGC responses, ignoring time. However, the linear reconstruction framework extends naturally to time-varying stimuli and RGC responses, and reconstructed image sequences. In a dynamic reconstruction, a given spike from a given RGC adds a specific spatio-temporal filter to the reconstruction, aligned in time with the spike (see Rieke et al, 1997). These contributions are summed over cells and over time. Although a quantitative assessment of performance and learning was not performed, dynamic reconstruction produced recognizable estimates of visual perception over time in both the natural and prosthesis simulation conditions (Fig. 10 and <u>supplemental movies</u>). In principle, these movies could be compared to patient reports with dynamic stimuli. Note that the dynamic reconstructions revealed persistent temporal structure in natural scenes, not visible in the still frames, which could improve performance on behavioral tasks.

**Figure 10:**
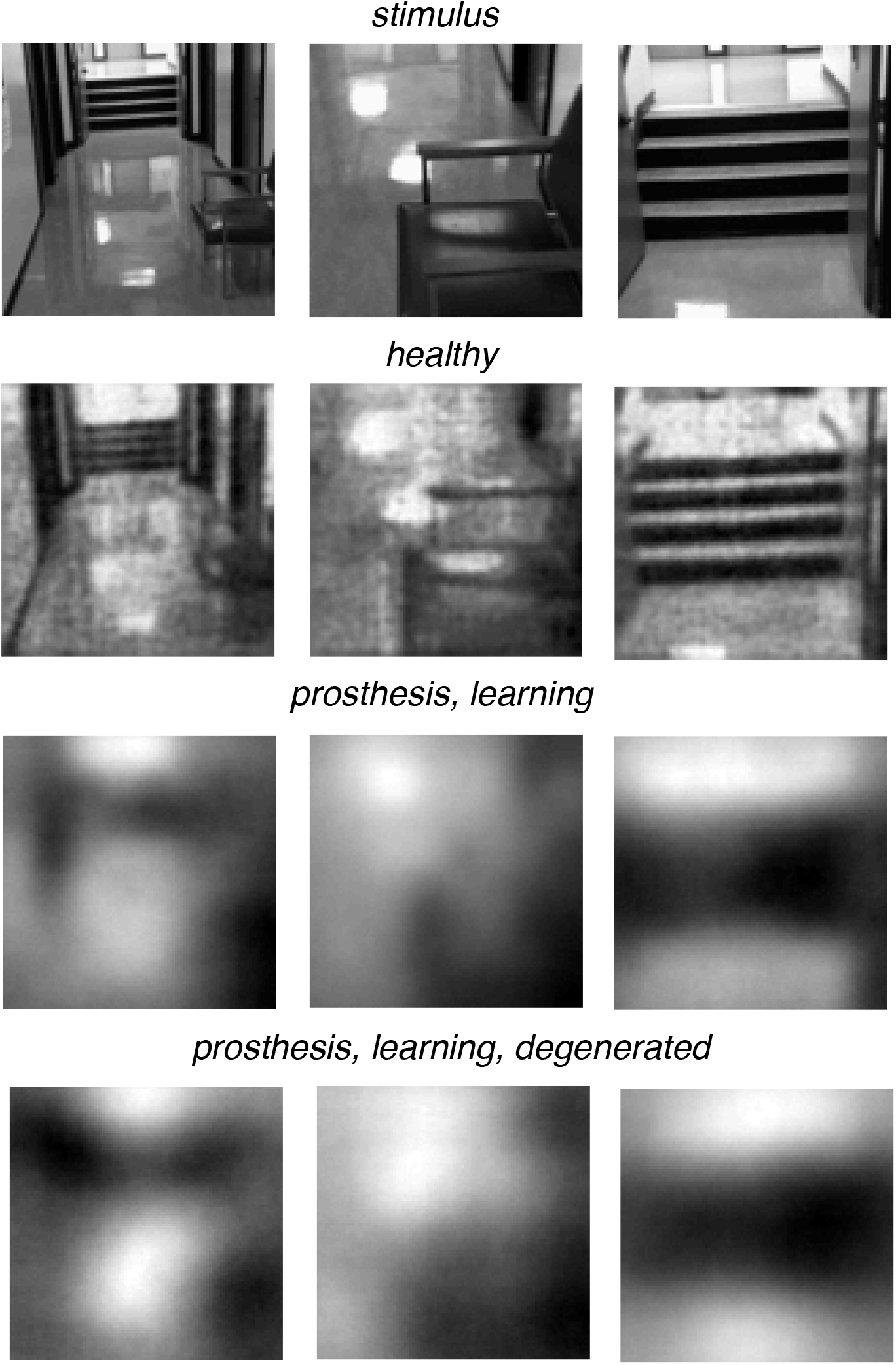
Individual frames from a dynamic movie stimulus (top row), reconstruction from the simulated healthy retinal response (second row), reconstruction from the prosthesis with learning (third row), and reconstruction from the prosthesis with learning with the degenerated retina.

## Discussion

A simulation framework was developed to explore visual perception and learning with a retinal prosthesis. The framework produced three primary predictions. First, as expected, artificial vision mediated by a prosthesis may be severely limited by the indiscriminate activation of ON and OFF cells, but dominance of ON cell activation could mitigate this effect. Second, learning by the patient could in principle produce substantial enhancements in performance, in part by compensating for the lack of cell type specificity. Finally, learning could enable somewhat greater benefits from technology enhancement, such as reduction of electrode spacing and current spread. Below, we discuss the assumptions underlying the simulation framework, its utility for future technology development, and major caveats.

### Impact of indiscriminate cellular activation

In healthy vision, roughly 20 different RGC types communicate different visual information to distinct targets in the brain, via their unique temporal patterns of activity (see Gollisch et al. 2010; Field and Chichilnisky 2007). Therefore, indiscriminate activation of different RGC types by a prosthesis produces a highly unnatural retinal code, potentially limiting the quality of artificial vision. The most obvious example is simultaneous activation of ON and OFF cells, which may produce conflicting visual signals in the brain. Although this issue is well known, its impact on visual perception has not been predicted quantitatively. The reconstruction approach provides a framework for measuring and interpreting the magnitude of this effect. For instance, a naive simulation in which ON and OFF cells respond equally to prosthesis activation, and their contributions to reconstruction are optimized for natural vision, predicts that artificial vision will be highly degraded. However, previous work suggests that either ON or OFF cells may respond more strongly to electrical stimulation (Werginz et al. 2015; Cottaris and Elfar 2005), and patient reports indicate that “bright” (rather than “dark”) phosphenes dominate increasingly over time (Stingl et al. 2013). A simple simulation of ON dominance – a linear reconstruction in which OFF cells are not used – revealed that it could substantially improve artificial vision. In principle, to obtain a quantitative simulation of ON dominance, patient performance in visual tasks (e.g. detection of increments or decrements) might be used to estimate a weighting factor between 0 and 1 applied to the OFF cell contribution. This factor could then be used as a coefficient in the linear reconstruction to make more specific predictions about patient percepts. The reconstruction framework may also be useful for assessing the value of developing retinal prostheses that can selectively stimulate distinct RGC types (Jepson et al. 2013, 2014, Nirenberg and Pandarinath 2012).

### Impact of learning

A closely related issue is the impact of learning by the patient, that is, changes in how the brain processes the retinal signal over time in the presence of retinal prosthesis stimulation. Although it is clear that learning occurs and improves artificial vision to some degree (Stingl et al. 2015; da Cruz et al. 2016), its precise impact and potential to improve artificial vision are unclear. The reconstruction approach provides a framework for predicting and interpreting the potential effects of learning. In a simulation of ideal (complete) learning, linear reconstruction optimized for prosthesis stimulation of the retina produced much more accurate estimates of the visual image, compensating to a substantial degree for the indiscriminate activation of ON and OFF cells. In principle, to obtain a more quantitative estimate of learning in the clinic, patient performance on visual tasks, compared to model performance, could be used to estimate a value between 0 and 1 indicating the degree of learning, that is, the relative contribution of the reconstruction filters optimized for prosthesis vs. healthy conditions. For example, a patient exhibiting a threshold of 0.3° on the Landolt C discrimination task (Fig. 9), roughly halfway between the learning and no learning conditions on a logarithmic scale, might be assigned a learning index of 0.5. This approach could make it possible to assess the improvements in performance over time in each patient relative to an upper bound set by “ideal” learning. Note that this interpretation assumes that the learning process involves enhanced perception, rather than behavioral strategies to deal with degraded perception in specific tasks. The choice of measurement of patient performance would be essential for distinguishing these forms of learning. Finally, note that the mechanisms by which this learning could take place are not addressed; instead, we provide a bound on the degree to which ideal learning could improve perception.

### Linearity, error measure, natural images

Three aspects of reconstruction were central to the implementation: the use of linear reconstruction, the choice of a squared error measure in computing reconstruction filters, and the optimization over natural images rather than another stimulus ensemble. Among reconstruction approaches, linear is the simplest, but others may exhibit higher performance. Nonlinear reconstruction is a large domain of inquiry, and improvements are already being made using artificial neural networks (Parthasarathy et al. 2017). The choice of pixel-wise squared error was made for technical convenience. A more perceptually relevant error measure might emphasize certain spatial scales, or specific image features, or perceptual decisions in the natural environment (Hoffman and Singh, 2012). A simple extension of the current approach would be to optimize reconstruction using perceptual image metrics, such as SSIM (Wang et al. 2004). Finally, the use of natural images is predicated on the idea that the visual system evolved to encode the structure in the natural environment. This assumption is important because the reconstruction filters are shaped by the statistics of visual inputs. However, the statistics of natural images are incompletely understood, and the particular statistical features which are most important to human evolution are even less clear. For example, human faces may be of high importance, but they represent only a tiny subset of the collection of images used in the simulation. This problem could potentially be explored by using more specific image databases (Parthasarathy et al. 2017).

### Related approaches

Most previous attempts to simulate the perceptual experience of patients with retinal prostheses have used a simple “scoreboard model” of phosphenes, in which pixels of an image are translated into perceived dots of light according to the layout of electrodes (Chen et al. 2009). A notable exception is a recent simulation framework that takes into account different electrical sensitivities of bipolar cells, RGCs somas, and RGC axons (Beyeler et al. 2017). This work reproduces clinical data and accounts for the perceptual effects of axon activation, temporal desensitization, and the distinct effects of pulse frequency and amplitude on phosphene attributes. However, the emphasis is quite different from the current work. Specifically, it is based on an empirical framework rather than a theoretical foundation such as optimal stimulus reconstruction, it does not address the indiscriminate activation of different retinal cell types, and it does not provide a prediction of the potential effects of learning. A comparison or combination of the two approaches in future patient data may be valuable.

### Image processing to optimize artificial vision

A potential enhancement of existing and future prosthesis technology is image processing prior to electrical stimulation, which could emphasize features of the visual scene of greatest significance to the patient, or more generally could produce a pattern of activation that is most likely to lead to accurate reconstruction. The rapid advance of cheap, low-power digital signal processing makes this kind of advance much more feasible. Although simple ideas such as edge enhancement have obvious potential, there is currently no framework within which to systematically test their impact in simulation. The framework proposed here could be used to optimize pre-processing of the image. By evaluating patient performance on specific tasks (Fig. 9), and comparing to the predictions, the impact of pre-processing could then be assessed and fine-tuned.

### Dynamic responses and reconstruction

The present study primarily tackled the case of static images, for simplicity. A more complete understanding would require analyzing dynamic vision, with a changing visual environment and head and eye movements. The methods presented here extend naturally to the dynamic case, by associating with each cell a spatiotemporal reconstruction filter, and summing those filters over cells and time. The results yield interpretable dynamic reconstructions (Fig. 10 and <u>supplemental movies</u>). In future work, some features of this simulation would need further investigation. First, although nonlinear reconstruction (see above) is readily accomplished with static images, this has not yet been performed with dynamic scenes, and the dimensionality of the problem is challenging. Second, perceptual measures of reconstruction accuracy, to improve upon the squared error metric, are less well developed for dynamic stimuli. Finally, appropriate training and testing stimuli are difficult to obtain, because the full dynamic properties of vision are usually not present in the movie data sets that are readily available.

### Model of cellular activation

In the present approach, a simple model of bipolar cell stimulation was used to capture the expected activation produced by a particular prosthesis technology (see Methods). However, different models would be required for different kinds of devices (e.g. see Beyeler et al. 2017). Furthermore, different electrodes in the device could activate nearby cells differently, based on factors such as their physical apposition to the retina and local damage due to disease, factors not accounted for here but that could potentially be calibrated in the clinic. A more complete model would also capture the increased spontaneous cellular activity associated with degeneration (Margolis et al. 2008), which varies between cell types (Sekirnjak et al. 2011), and effectively introduces noise into the visual signal transmitted to the brain. The simple simulation of degeneration here provides a sense for the magnitude of the impact of such factors, but more exploration is needed. Variations can be naturally incorporated into the reconstruction framework presented here, by modification of the cellular activation module or retinal circuit module, without changing the underlying logic of the approach. The learning simulation could then be used to study the potential for patients to overcome the different kinds of cellular activation arising from these factors.

### Interpreting the visual experience of patients

A challenge associated with assessing prosthesis function is attempting to understand what patients see. Verbal reports are helpful, but it is extremely difficult to use the reports provided by untrained blind patients to obtain a complete sense of their visual experience. Because the reconstruction framework produces images as predictions of visual experience, it could be used by clinicians to compare with and interpret patient verbal reports. Such feedback may help to fine-tune the assumptions and parameters of the simulation (see Beyeler et al. 2017), such as the degree of ON dominance or learning (see above). Note that variations in performance that are relevant to the clinic, such as surgical approach, were not considered here.

### Assessing the potential of technological developments

The development of retinal prostheses is ongoing, and represents a major international effort (Goetz and Palanker 2016). Given the enormous resources required to develop new technologies and obtain regulatory approval, as well as the risks to human patients receiving the implants, simulations that can predict reasonably accurately the performance of envisioned future devices could be immensely valuable (Beyeler et al. 2017). The potential for such predictions is illustrated by examining the dependence of reconstruction on features such as electrode spacing and current spread (Figs. 6 and 7), which suggests that the biological limitations of the prosthesis need to be considered alongside the technical constraints.

### Caveats and future

Several other uncertain features of the approach deserve mention. First, the linear reconstruction concept underlying the approach is perhaps the most obvious first step, but it is difficult to test explicitly. Second, the model of retinal activation by the photovoltaic implant and the model of healthy retinal processing are both rudimentary, and will need to be improved to better capture temporal phenomena, nonlinearities, noise, and cell type specificity. Furthermore, it will be challenging to produce an accurate simulation of the retinal activity produced by a specific prosthesis technology implanted in a specific patient with a particular disease process and treatment history, because of variations in the electrical interface and the state of the patient’s central visual system. Finally, technical features of the reconstruction approach can surely be improved, as discussed above, and such improvements should be guided by patient reports and performance. Thus, the framework presented here is not intended to be used unmodified in a clinical application. Instead, it may be considered a proof of concept and starting point for future efforts.

## Acknowledgements

We thank Vincent Bismuth and Daniel Palanker for valuable discussions and comments on the manuscript, Nishal Shah for technical assistance and useful input, and Georges Goetz for helpful discussions.

